# TIMP-3 facilitates binding of target metalloproteinases to the endocytic receptor LRP-1 and promotes scavenging of MMP-1

**DOI:** 10.1101/2019.12.23.886762

**Authors:** Anna P. Carreca, Veronica Pravatà, Gillian Murphy, Hideaki Nagase, Linda Troeberg, Simone D. Scilabra

## Abstract

Matrix metalloproteinases (MMPs) and the related families of disintegrin metalloproteinases (ADAMs) and ADAMs with thrombospondin repeats (ADAMTSs) play a crucial role in the turnover of extracellular matrix (ECM) components and shedding of cell-surface molecules. The proteolytic activity of metalloproteinases is post-translationally regulated by their endogenous inhibitors, known as tissue inhibitors of metalloproteinases (TIMPs). Several MMPs, ADAMTSs and TIMPs have been reported to be endocytosed by the low-density lipoprotein receptor-related protein-1 (LRP-1). Different binding affinities of these proteins for the endocytic receptor correlate with different turnover rates which, together with differences in their mRNA expression, determines their nett extracellular levels.

In this study, we used surface plasmon resonance to evaluate the affinity between LRP-1 and a number of MMPs, ADAMs, ADAMTSs, TIMPs and metalloproteinase/TIMP complexes. This identified MMP-1 as a new LRP-1 ligand. Among the proteins analysed, TIMP-3 bound to LRP-1 with highest affinity (*K*_D_ = 1.68 nM). Additionally, we found that TIMP-3 can facilitate the clearance of its target metalloproteinases by bridging their binding to LRP-1. For example, the free form of MMP-1 was found to have a *K*_D_ of 34.6 nM for LRP-1, while the MMP-1/TIMP-3 complex had a 7-fold higher affinity (*K*_D_ = 4.96 nM) for the receptor. TIMP-3 similarly bridged binding of MMP-13 and MMP-14 to LRP-1. TIMP-1 and TIMP-2 were also found to increase the affinity of target metalloproteinases for LRP-1, albeit to a lesser extent. These findings suggest that LRP-1 scavenging of TIMP/metalloproteinase complexes may be a general mechanism by which inhibited metalloproteinases are removed from the extracellular environment.

## INTRODUCTION

Extracellular matrix (ECM) turnover is an important feature of several physiological processes, including development, organogenesis, wound healing and tissue remodelling. The matrix metalloproteinases (MMPs) are a major class of proteinases involved in ECM turnover (1). The related disintegrin metalloproteinases (ADAMs) and ADAMs with thrombospondin motifs (ADAMTSs) are also involved in ECM turnover by processing of cell surface molecules and various ECM components, respectively (2,3). Tissue inhibitors of metalloproteinases (TIMPs) are wedge-shaped proteins that regulate the activity of MMPs, ADAMs and ADAMTSs by interacting with the active site cleft of the enzymes and forming a tight complex with a 1:1 stoichiometry (4). Under physiological conditions, ECM turnover is regulated by the balance between metalloproteinases and TIMPs, and disruption of this balance is associated with pathological conditions characterised by enhanced ECM degradation, such as arthritis, cancer and cardiovascular disorders (5-7).

The expression of metalloproteinases and TIMPs can be regulated at different levels, ranging from regulation of their transcription by cytokines and growth factors, to post-transcriptional regulation by RNA interference. In addition, the catalytic activity of several metalloproteinases can be post-translationally activated by proteolytic removal of their inhibitory pro-domain (4,8). Moreover, a major mechanism that regulates the extracellular levels of several MMPs [including MMP-2 (9), MMP-9 (10) and MMP-13 (11,12)] ADAMTSs [including ADAMTS-1 (13), ADAMTS-4 (14) and ADAMTS-5 (15)] and TIMPs [including TIMP-1 (16), TIMP-2 (9), TIMP-3 (17,18)] is endocytosis and subsequent lysosomal degradation *via* the scavenger receptor low-density lipoprotein receptor-related protein 1 (LRP-1) (19,20). LRP-1 is considered to be a master regulator of ECM turnover, since the affinity of individual metalloproteinases and TIMPs for LRP-1 determines their extracellular half-life and so controls the balance between ECM degradation and deposition. In this study, we used surface plasmon resonance (SPR) to carry out a systematic analysis of the affinity of several MMPs, ADAMs, ADAMTSs, TIMPs and metalloproteinase/TIMP complexes for LRP-1. This allowed us to investigate the hierarchy of LRP-1 ligands within the metalloproteinase activity network.

Among all proteins analysed, we found that TIMP-3 bound to LRP-1 with the highest affinity. Furthermore, TIMP-3 was able to promote scavenging of its target enzymes, with TIMP-3/metalloproteinase complexes having lower *K*_D_ values for LRP-1 than observed for the enzymes alone. *K*_D_ values for TIMP-3/metalloproteinase complexes were almost indistinguishable from that of TIMP-3 alone, indicating that the inhibitor can bridge the binding of metalloproteinases to LRP-1. TIMP-1 and TIMP-2 were also able to promote scavenging of their target enzymes, albeit to a lesser extent, indicating that TIMP-mediated scavenging may be a general mechanism by which inhibited metalloproteinases are cleared from the extracellular environment.

## EXPERIMENTAL PROCEDURES

### Materials and cells culture

HTB94 human chondrosarcoma cells were from American Culture Type Collection (Manassas, VA, USA), and maintained in DMEM with 10 % FCS, 100 U/ml penicillin and 100 U/ml streptomycin at 37 °C in 5 % CO_2_. Dulbecco’s modified Eagle’s medium (DMEM), L-glutamine, penicillin/streptomycin, fetal calf serum (FCS), hygromycin B and trypsin-EDTA from PAA Laboratories (Somerset, UK); DMEM without L-glutamine, cysteine, methionine or cystine from MP Biomedicals (Solon, OH, USA). BSA was from Sigma-Aldrich (Dorset, UK); LRP-1 was from Biomac (Leipzig, Germany); ADAM10 and ADAM17 ectodomain were from R&D Systems (Abingdon, UK). The catalytic domain of human MMP-1 (MMP-1ΔC), full length MMP-1 and MMP-1 E200A (21); recombinant His-tagged RAP (15); FLAG-tagged ADAMTS-4 lacking the C-terminal spacer domain (22); FLAG-tagged ADAMTS-5 lacking the C-terminal thrombospondin domain and lacking the C-terminal thrombospondin and spacer domains (23); FLAG-tagged TIMP-3 and FLAG-tagged N-TIMP-3 (24); TIMP-1 and TIMP-2 (25); MMP-3 catalytic domain (MMP-3ΔC) (26); MMP-9 (27); MMP-13 E204A (28) were prepared as previously described. The ectodomain of MMP-14 was expressed in *E. coli* BL21(DE3). Inclusion bodies were isolated and refolded as described by Huang et al. (29).

### Surface Plasmon Resonance analysis

The affinity of various ligands to LRP-1 was evaluated using a BIAcore T200 (GE, Amersham, UK). LRP-1 was immobilized on a CM5 sensor chip by amine coupling using N-hydroxysuccinimide (NHS), according to the manufacturer’s instructions. Immobilization was performed at 10 μl/min until reaching a target ligand concentration of 3000 RUs on the chip. The instrument was maintained at 25 °C. SPR analysis was performed in 150 mM NaCl, 10 mM CaCl2, 50 mM Tris-HCl buffer, 0.01 % Tween-20, pH 7.5, at a flow rate of 30 μl/min. After each cycle, the sensor chip was regenerated by injecting 30 μl of 10 mM glycine-HCl buffer, pH 2.5. All ligands were injected over the immobilized LRP-1 at 6 concentrations ranging from 0 to 80 nM (except BSA which was used as a negative control and injected at up to 2 μM; MMP-1ΔC that was injected at up to 2 μM; and MMP-14, ADAM-10 and ADAM-17 ectodomain that were injected at up to 160 nM). Metalloproteinase/TIMP complexes were pre-formed *in vitro* by incubating equimolar concentrations of enzyme and inhibitor for 1 h at 37 °C. Complexes were injected at 5 different concentrations (5 nM, 10 nM, 20 nM, 40 nM and 80 nM). The *K*_i_ of MMP-1 for TIMP-1, TIMP-2 and TIMP-3 is 0.38 nM, 1.03 nM and 1.1 nM respectively (17,22), therefore metalloproteinase/TIMP complexes should be stable at all the concentrations used for injection. Each determination was performed in triplicate. BIAevaluation software was used to analyze the resulting sensorgrams and determine the association (*k*_a_) and dissociation (*k*_d_) rate constants, and the constant of dissociation (*K*_D_). Binding data were fitted in a one-site binding model.

### Expression and purification of [^35^S]TIMP-3 and [^35^S]MMP-1

[^35^S]TIMP-3 was prepared by metabolic labelling as previously described (30). Human proMMP-1 was cloned into pCEP4 expression plasmid with a FLAG-tag between the signal and prodomain. [^35^S]MMP-1 was generated by a modification of the protocol for [^35^S]TIMP-3 (30). HEK-293/EBNA cells transfected with proMMP-1/pCEP4 were grown to confluence in a 150 cm^2^ flask, washed once and starved for 2 h in serum-free DMEM without L-glutamine, cysteine, methionine or cysteine. Cells were then grown for a further 4 days in serum-free DMEM without L-glutamine, cysteine, methionine or cysteine, and supplemented with [^35^S]Met/[^35^S]Cys (500 μCi of Redivue Pro-Mix L-[^35^S] *in vitro* Cell Labelling Mix, GE Healthcare, Buckinghamshire, UK). Conditioned media were collected after 4 days, centrifuged to remove cell debris and applied to a 2 ml anti-FLAG M2-agarose column (Sigma-Aldrich, Dorset, UK). The resin was washed extensively in TNC buffer [50 mM Tris-HCl (pH 7.5), 150 mM NaCl, 10 mM CaCl_2_]. Bound [^35^S]proMMP-1 was eluted with 200 μg/ml FLAG peptide (Sigma-Aldrich, Dorset, UK). After elution with FLAG peptide, [^35^S]proMMP-1 was activated with APMA as previously described (31). Cleavage of the pro-domain was confirmed by SDS-PAGE, and the pro-domain was removed using a PD10 desalting column (GE, Amersham, UK). The active concentration of [^35^S]MMP-1 was determined by titration against a known concentration of the N-terminal domain of TIMP-3 (N-TIMP-3).

### Endocytosis of [^35^S]TIMP-3, [^35^S]MMP-1 and MMP-1/TIMP-3 complexes

HTB94 cells were plated at a density of 1×10^6^ cells/well, rested overnight and then washed with serum-free DMEM. Cells were then incubated for up to 24 h with either 10 nM [^35^S]TIMP-3 or 10 nM [^35^S]TIMP-3/enzyme complexes, preformed *in vitro* by incubating [^35^S]TIMP-3 and enzyme for 1 h at 37 °C. At specified times, the conditioned media were removed and 5 % (v/v) trichloroacetic acid (TCA) added (4 °C, overnight). The TCA-soluble fraction (2 ml) was separated from the TCA-insoluble fraction by centrifugation (13000 rpm, 15 min, 4 °C). The TCA-insoluble pellet was dissolved in 1 N NaOH (500 μl). The cell layer was washed with ice-cold PBS and solubilized in 1 N NaOH [1 ml, 1 h, room-temperature (RT)]. Each fraction was mixed with 3 ml of scintillant (Goldstar Multipurpose Liquid Scintillation Cocktail, Meridian, Surrey, UK) and radioactivity counted. TCA-soluble radioactivity present in the [^35^S]TIMP-3 preparation prior to incubation with cells (5-15 % of total radioactivity) was subtracted from the amount of TCA-soluble radioactivity at each time point. The amount of radioactivity in the different fractions was calculated as a percentage of the total amount of [^35^S]TIMP-3 radioactivity added to the cells. Endocytosis of [^35^S]MMP-1 and [^35^S]MMP-1/TIMP complexes was evaluated in a similar manner.

## RESULTS

### TIMP-3 bridges the binding of metalloproteinases to LRP-1 *in vitro*

We have previously shown that TIMP-3 directly binds to LRP-1 using ELISA (30). In order to understand this interaction in greater detail, we used surface plasmon resonance to determine the affinity of TIMP-3 for LRP-1 immobilised on a BIAcore sensor chip. Receptor-associated protein (RAP), which is a well-established LRP-1 ligand (32,33), was used as positive control for binding. We measured a *K*_D_ of 0.6 nM (Table 1), in good agreement with the previous determination of Lee and colleagues, who reported a *K*_D_ of 1.4 nM for this interaction (34). The negative control, BSA, did not show any binding at concentrations up to 2 μM. TIMP-3 bound to LRP-1 with high affinity (*K*_D_ = 1.68 nM, Table 1, Figure 1A). The N-terminal domain of TIMP-3 (N-TIMP-3) bound to LRP-1 with a *K*_D_ of 5.65 nM (Table 1), confirming that the minimal binding determinants reside in the N-terminal domain of the inhibitor. This is in line our previous finding that N-TIMP-3 is endocytosed by chondrosarcoma cells in an LRP-1-dependent manner with similar kinetics to that of TIMP-3 (30). TIMP-1 and TIMP-2 also bound to LRP-1, albeit with 18-fold and 16-fold lower affinities respectively (Table 1).

**Table 1.**
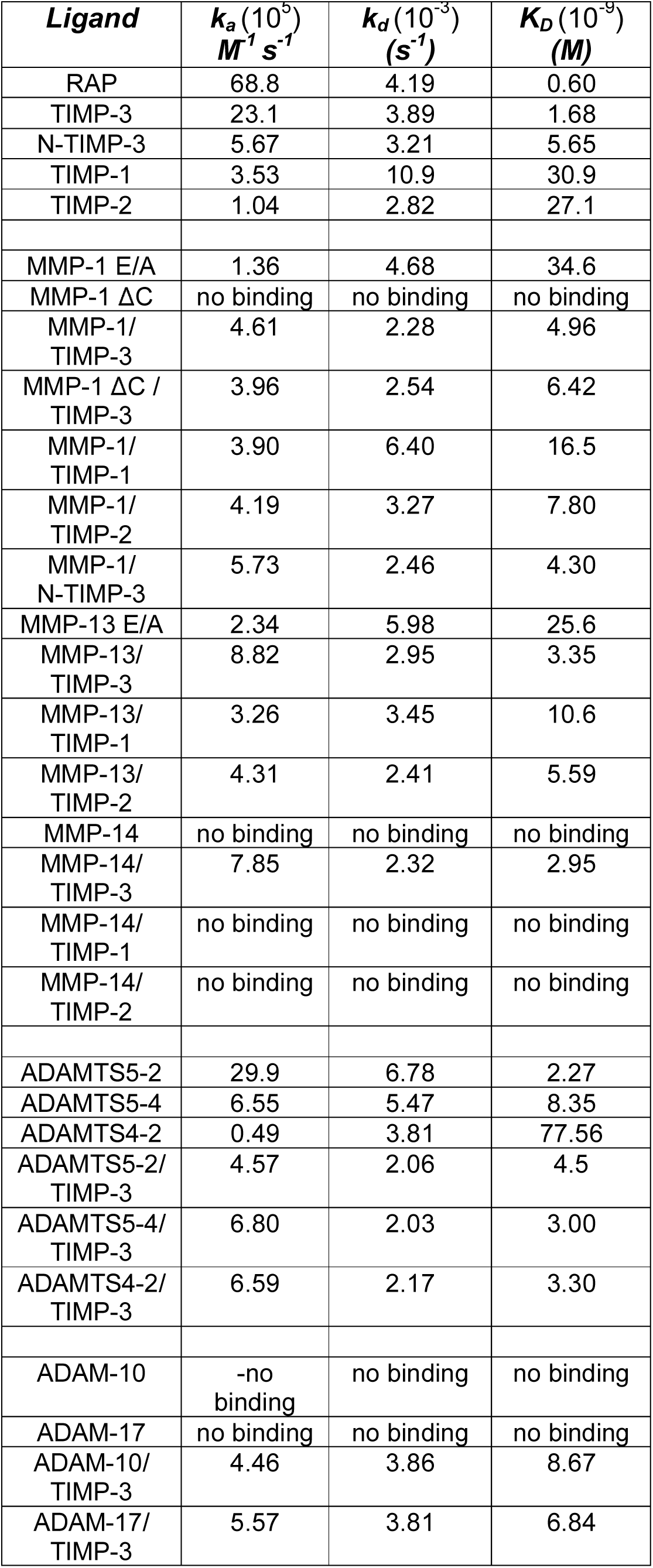
Kinetics of ligand binding to LRP-1 measured by SPR. LRP-1 was immobilized on a CM5 sensor chip and the binding constant of different analytes were determined. Each determination was performed in triplicate at six concentration for each analyte [0-80 nM except ADAMTS4-2 (0-180 nM), BSA, MMP-1 ΔC and MMP-14 ectodomain (0-2 μM)]. Binding constants were calculated using the BIOevaluation software.

**Figure 1.**
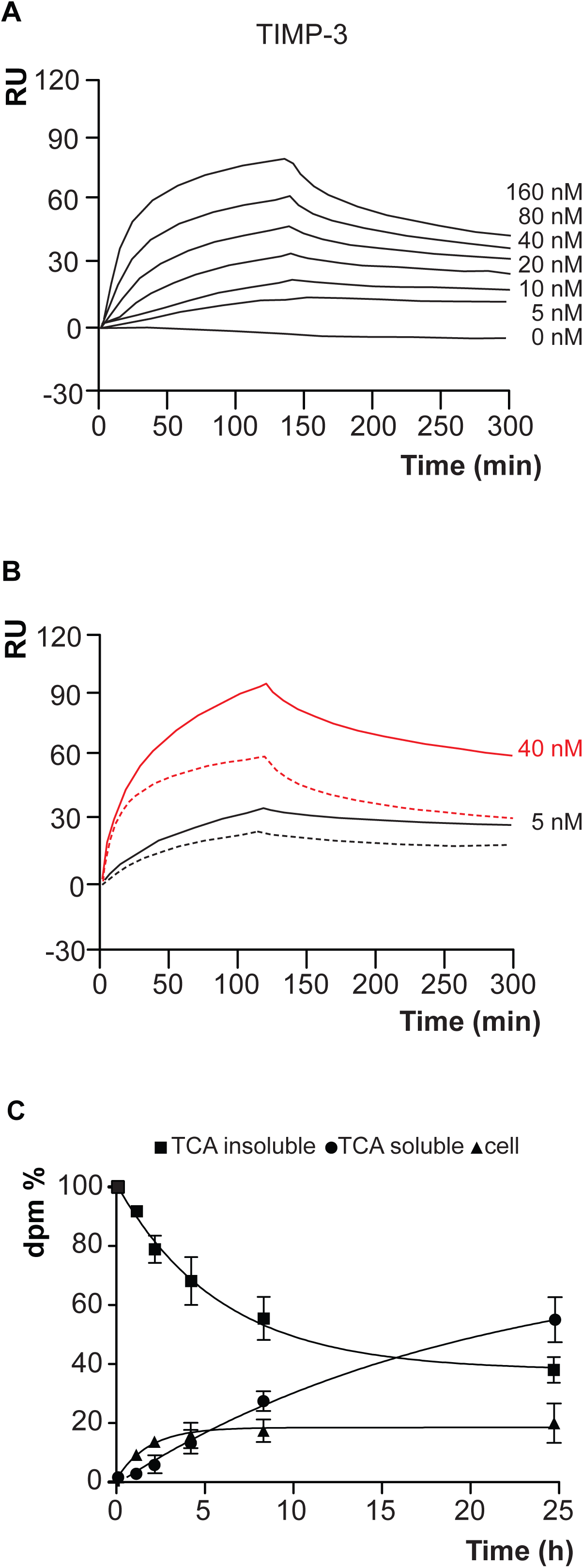
Binding of TIMP-3 to LRP-1 and its internalisation by HTB94 cells. **A**. Sensograms from a surface plasmon resonance analysis of the interaction between immobilized LRP-1 and TIMP-3, passed over a chip at six concentrations (0-80 nM). **B**. Sensograms showing the binding of TIMP-3 (solid line) and MMP-1/TIMP-3 complex (dashed line) to immobilized LRP-1 at 10 nM (black lines) and 40 nM (grey lines). **C**. [^35^S]TIMP-3 (10 nM) added to HTB94 chondrosarcoma cells, and radioactivity in different cell fractions monitored over time (n = 3).

#### MMP-1 is a new LRP-1 ligand

Because MMP-1 undergoes autocatalysis at room temperature, we used a catalytically inactive form of MMP-1 [MMP-1 (E200A)] to investigate binding of this enzyme to LRP-1. SPR analysis showed that MMP-1 (E200A) bound to LRP-1 with a *K*_D_ of 34.6 nM (Table 1). The MMP-1 catalytic domain (MMP-1ΔC) showed no binding to LRP-1 even at micromolar concentrations (Table 1), indicating that the hemopexin domain is required for binding of MMP-1 to the receptor. This is in agreement with previous studies showing that the hemopexin domain of MMP-9 mediates the majority of MMP-9 interaction with LRP-1 (35).

#### TIMPs facilitate binding of MMP-1 and MMP-13 to LRP-1 in vitro

Emonard *et al*. reported that the pro-MMP-2/TIMP-2 complex has a higher affinity for LRP-1 than either pro-MMP-2 or TIMP-2 alone (20). We thus investigated whether interaction with TIMP-1, TIMP-2 or TIMP-3 could increase MMP-1 binding to LRP-1. MMP-1/TIMP complexes were made *in vitro* by incubating equimolar concentrations of MMP-1 and TIMPs (5-80 nM) for 1 hour at 37 °C before addition to immobilised LRP-1. Previously reported *K*_i_ values for MMP-1 inhibition by TIMP-1, TIMP-2 and TIMP-3 are 0.38 nM, 1.03 nM and 1.1 nM respectively, indicating that MMP-1/TIMP complexes remain stable at the concentrations used (17,22). Interaction with TIMP-3 increased MMP-1 affinity for LRP-1 by 7-fold, to a *K*_D_ of 4.96 nM (Table 1). SPR quantifies molecular interactions by detecting changes in refractive index in the aqueous layer close to the sensor chip surface (36). Binding of large complexes is associated with greater changes in refractive index. Figure 1B shows MMP-1/TIMP-3 complexes induce a higher response than the same concentration of TIMP-3 alone, indicating that under these experimental conditions, MMP-1 and TIMP-3 bind to LRP-1 as a complex. MMP-1ΔC had no detectable affinity for LRP-1 on its own, but when in complex with TIMP-3, a *K*_D_ of 6.42 nM was detected (Table 1), indicating that TIMP-3 is largely responsible for the binding of this complex to LRP-1. Interaction with TIMP-1 and TIMP-2 also increased MMP-1 affinity for LRP-1, by 2- and 4.5-fold respectively. Unlike for TIMP-3, the affinities of the complexes are higher than that for TIMP-1 or TIMP-2 alone (Table 1), suggesting that MMP-1 either contributes to the binding, or that it stabilises a TIMP-1/-2 conformation with increased LRP-1 affinity.

TIMPs were also able to promote binding of other MMPs to LRP-1. To study MMP-13, which undergoes autodegradation at room temperature, we used a catalytically inactive mutant, namely MMP-13 (E204A). This collagenase had a *K*_D_ value of 25.6 nM for LRP-1 (Table 1). Interaction with TIMP-3 increased its affinity for LRP-1 by 7.6-fold, while interaction with TIMP-1 or TIMP-2 increased its affinity by 2.4- and 4.6-fold respectively. (Table 1). The ectodomain of MMP-14 showed no detectable binding to LRP-1 even at micromolar concentrations. Incubation with TIMP-3 facilitated binding with a *K*_D_ of 2.95 nM, while incubation with TIMP-1 or TIMP-2 did not detectably improve affinity (Table 1).

#### TIMP-3 increases the affinity of ADAMTS-4 and deletion mutants of ADAMTS-5 for LRP-1

The aggrecanase ADAMTS-5 is an LRP-1 ligand, with the LRP-1-binding determinants localized in the ancillary domains of the enzyme. Affinity for LRP-1 thus decreases with consecutive deletion of the non-catalytic C-terminal domains (14,37). Here, we found that ADAMTS-5 lacking the C-terminal thrombospondin domain (ADAMTS5-2) bound effectively to LRP-1, with a *K*_D_ of 2.27 nM. Deletion of the cysteine-rich and spacer domain (ADAMTS5-4) reduced the affinity of ADAMTS-5 for LRP-1 to 8.35 nM. Binding to TIMP-3 had little effect on ADAMTS5-2 affinity for LRP-1 (*K*_D_ increased to 4.5 nM), but improved ADAMTS5-4 affinity for LRP-1 (*K*_D_ = 3.00 nM) (Table 1).

ADAMTS-4 had lower affinity for LRP-1 (*K*_D_ = 77.56 nM) than ADAMTS-5 (*K*_D_ = 2-8 nM) (Table 1). Binding to TIMP-3 improved the *K*_D_ to 3.3 nM, indicating that TIMP-3 can also mediate ADAMTS-4 scavenging.

#### TIMP-3 also improved affinity of ADAM10 and ADAM17 for LRP-1

Binding of LRP-1 was undetectable for the free form of either ADAM10 or ADAM17, while binding to TIMP-3 improved their affinity markedly, with *K*_D_ values of 8.67 for ADAM10/TIMP-3 and 6.84 for ADAM17/TIMP-3. This suggests that TIMP-3 can facilitate the binding of these enzymes to LRP-1. In conclusion, a systematic SPR analysis showed that TIMP-3 binds to LRP-1 with the highest affinity among TIMPs. TIMPs generally increased affinity of the binding of specific collagenases (MMP-1 and −13) to LRP-1, with TIMP-3 mediating this process to the greatest extent.

### Complex formation with metalloproteinases does not affect the kinetics of TIMP-3 endocytosis

TIMP-3 is rapidly internalised by various cell types via LRP-1. [^35^S]radiolabeling of TIMP-3 and subsequent detection of radioactivity in different cell compartments has been used to follow TIMP-3 endocytosis (18). We used this method to investigate whether complex formation with metalloproteinases had an effect on the rate of [^35^S]TIMP-3 endocytosis. [^35^S]TIMP-3/metalloproteinase complexes were generated *in vitro* by incubating 10 nM [^35^S]TIMP-3 with an equimolar concentration of various metalloproteinases for 1 h at 37 °C. Formation of 1:1 stoichiometric complexes and their stability over 24 h were confirmed by measuring their activity against specific synthetic substrates. As previously shown (18), levels of [^35^S]TIMP-3 in the TCA insoluble fraction of the conditioned media (containing intact not yet internalised [^35^S]TIMP-3) decreased over time, while cell-associated [^35^S]TIMP-3 ([^35^S]TIMP-3 inside the cells after internalisation) and [^35^S]TIMP-3 in the TCA soluble fraction of the conditioned media ([^35^S]TIMP-3 released into the media after internalisation and lysosomal degradation) increased over time (Figure 1C). [^35^S]TIMP-3/MMP-1 complex was internalised with indistinguishable kinetics compared to [^35^S]TIMP-3 alone (Figure 2A). Similar results were obtained when [^35^S]TIMP-3 was in complex with MMP-1ΔC, which had no affinity at all for LRP-1 *in vitro* (Figure 2A). In addition to full length MMP-1 and MMP-1ΔC, clearance of [^35^S]TIMP-3 was not affected when the inhibitor was in complex with MMP-3ΔC, MMP-9 or MMP-14 ectodomain (Figure 2B). K_D_ of TIMP-3 for LRP-1 when in complex with ADAMTS5-2 or ADAMTS4-2 was similar to that of TIMP-3 alone. In agreement, complex formation with ADAMTS5-2 or ADAMTS4-2 did not significantly affect the internalization kinetics and clearance of [^35^S]TIMP-3 (Figure 2C-D).

**Figure 2.**
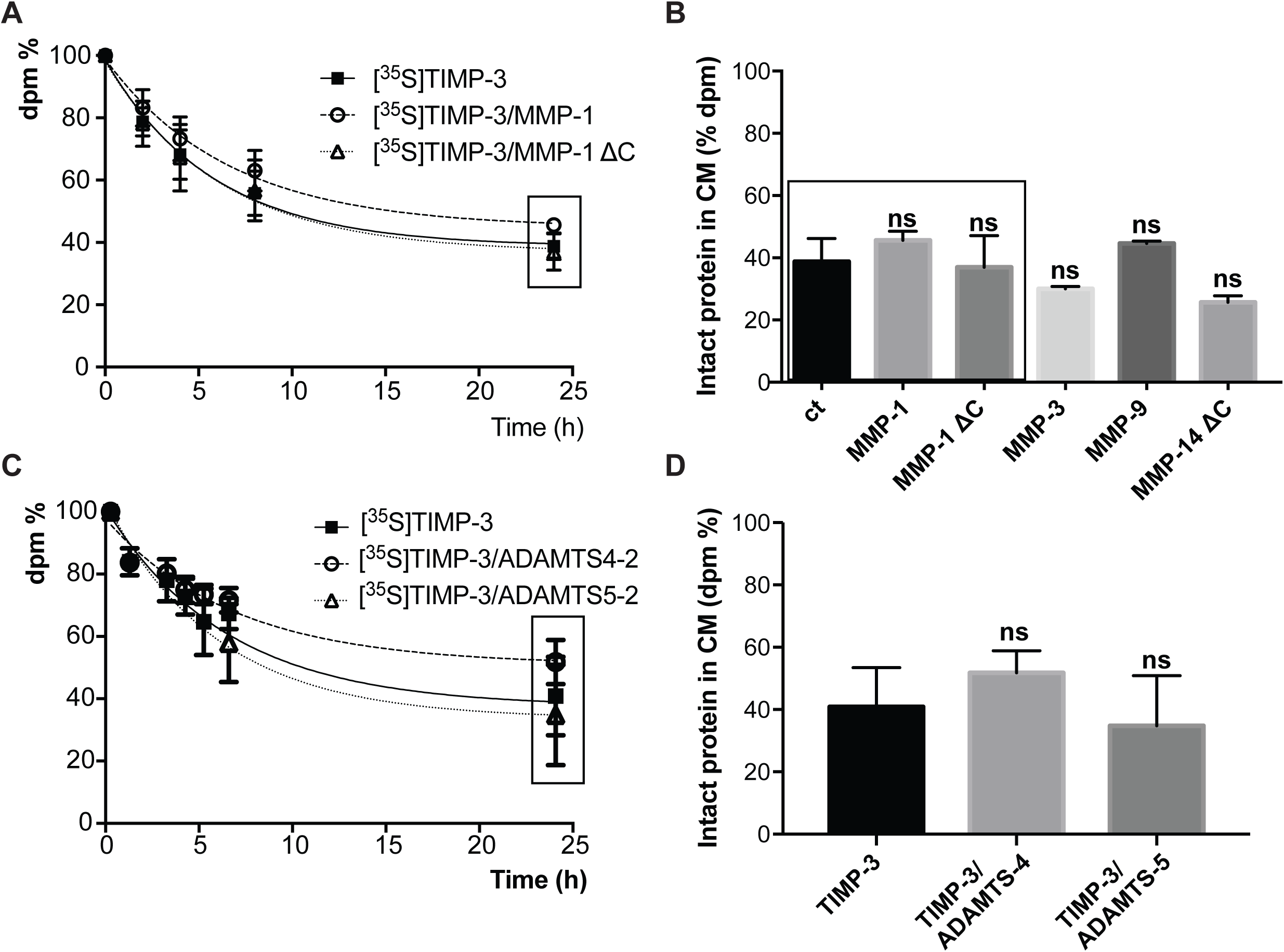
Internalisation of TIMP-3 in complex with a number of MMPs and ADAMTSs. [^35^S]TIMP-3 was incubated with either MMP-1, MMP-1ΔC, MMP-3, MMP-9, MMP-14, ADAMTS4-2 or ADAMTS5-2 for 1 h at 37 °C to make a complex *in vitro*. **A**. HTB94 cells were incubated with either [^35^S]TIMP-3 alone (solid squares, solid black line) or [^35^S]TIMP-3/MMP-1 complex (open circles, black dashed line) or [^35^S]TIMP-3/MMP-1ΔC complex (open triangles, dotted black line). At indicated time points intact [^35^S]TIMP-3 in the conditioned media was measured. **B**. Clearance of [^35^S]TIMP-3 after 24 h incubation with HTB94 cells was evaluated for the inhibitor alone or in complex with MMP-1, MMP-1ΔC MMP-3, MMP-9 and MMP-14 catalytic domain and shown as mean values of intact [^35^S]TIMP-3 remained in the media□±□standard deviation (analysed by one-way ANOVA, followed by Dunnett’s multiple comparisons test where [^35^S]TIMP-3 was taken as a control; ns non significant; n=3). Values for [^35^S]TIMP-3, [^35^S]TIMP-3/MMP-1 and [^35^S]TIMP-3/MMP-1ΔC were extrapolated from panel A (symbols inside the black box). **C**. Similar analysis was performed for [^35^S]TIMP-3 alone (solid squares, black solid line) or [^35^S]TIMP-3/ADAMTS4-2 complex (black dashed line and open circles) or [^35^S]TIMP-3/ADAMTS5-2 (black dotted line and open triangles). **D**. Clearance of [^35^S]TIMP-3 alone [^35^S]TIMP-3/ADAMTS4-2 or [^35^S]TIMP-3/ADAMTS5-2 after 24 h was extrapolated from panel C (symbols inside the black box), and represented as mean values of intact [^35^S]TIMP-3 remained in the media □±□ standard deviation and analysed by one-way ANOVA, followed by Dunnett’s multiple comparisons test where [^35^S]TIMP-3 was taken as a control; ns non significant; n=3).

In conclusion, the internalisation rate of TIMP-3 is not affected by complex formation with metalloproteinases, suggesting that the inhibitor can drive the internalization of TIMP-3/metalloproteinase complexes.

### MMP-1 is endocytosed by cells and TIMP-3, but not other TIMPs, increases its endocytosis rate

SPR analysis showed that MMP-1 directly binds to LRP-1 through its hemopexin domain, suggesting that it could be internalised in an LRP-1-dependent manner, similarly to other MMPs, including MMP-2, MMP-9 and MMP-13 (9,10,12). Thus, in order to investigate whether MMP-1 could be internalised in a cellular system, it was metabolically radiolabelled with [^35^S], purified and 10 nM [^35^S]MMP-1 incubated with HTB94 cells. Levels of intact [^35^S]MMP-1 in the conditioned media decreased of about 20% over 24h, while [^35^S]MMP-1 inside the cells and in the TCA soluble fraction ([^35^S]MMP-1 fragments deriving from lysosomal degradation and released in the conditioned media) increased over the time, indicating that MMP-1 is indeed internalised by cells, although at lower rate compared to TIMP-3 and in agreement with their different K_D_ values for LRP-1 (Figure 3A and 3E). Conversely, when 10 nM [^35^S]MMP-1 was pre-incubated with an equimolar concentration of TIMP-3 to form a [^35^S]MMP-1/TIMP-3 complex, its internalisation increased, showing endocytosis kinetics similar to that of [^35^S]TIMP-3 alone (Figure 3B and 3E). Neither TIMP-1 nor TIMP-2 promoted [^35^S]MMP-1 endocytosis under the experimental conditions (Figure 3C-E). Indeed, at concentrations used in this assay, MMP-1/TIMP-1 and MMP-1/TIMP-2 complexes had reduced binding to LRP-1 on SPR compared to an equal concentration of MMP-1/TIMP-3 complexes (Figure 3F).

**Figure 3.**
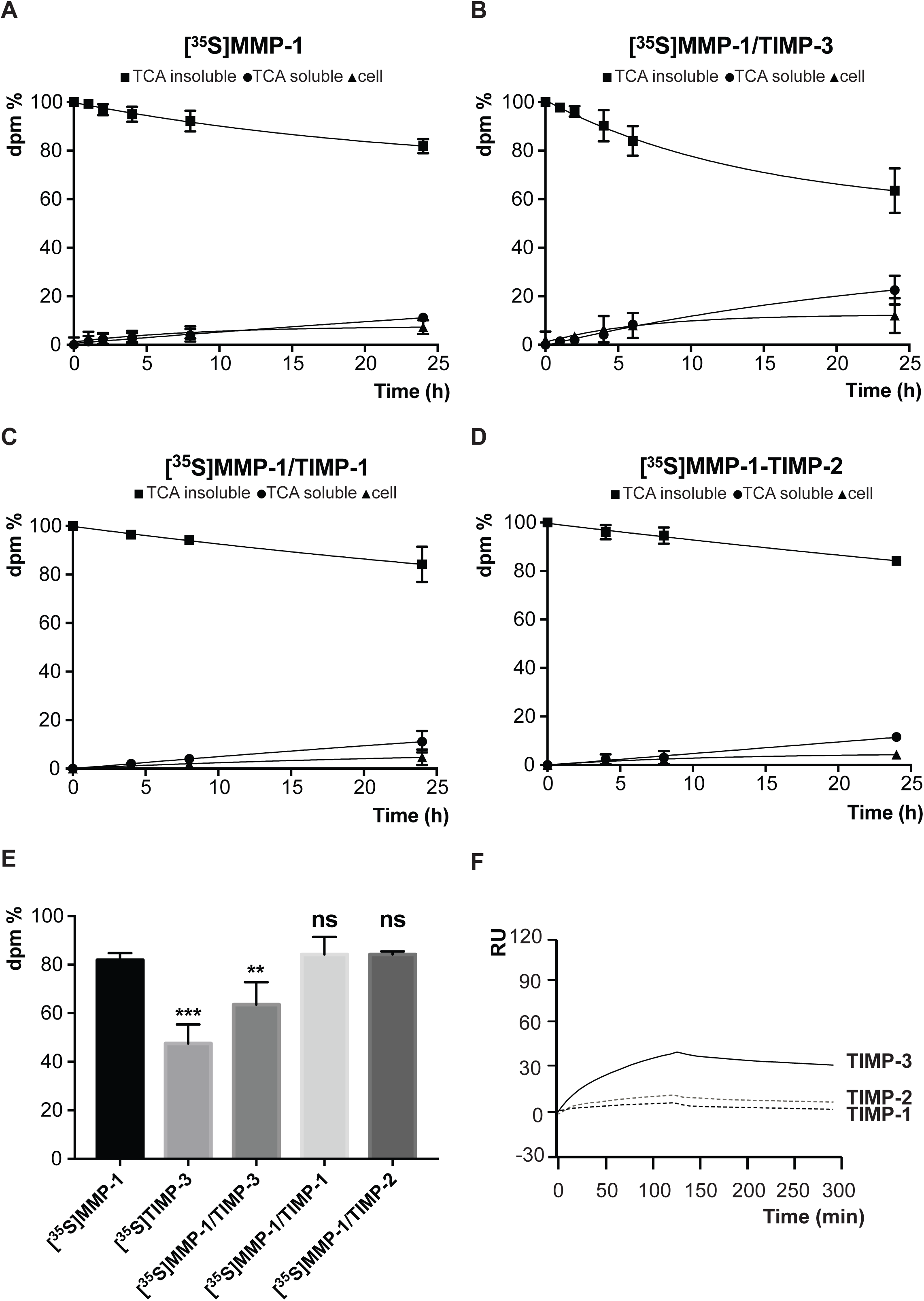
MMP-1 is internalised by cells and its uptake is increased by formation of a complex with TIMP-3. A-D HTB94 cells were incubated with 10 nM [^35^S]MMP-1 or with pre-formed [^35^S]MMP-1/TIMP-3, [^35^S]MMP-1/TIMP-1 or [^35^S]MMP-1/TIMP-2 complexes. At indicated times, the TCA-precipitated amount of intact [^35^S]MMP-1 (TCA-insoluble fraction, squares) and the degraded [^35^S]MMP-1 (TCA-soluble fraction, circles) in the conditioned media, together with the cell-associated [^35^S]MMP-1 (triangles) were measured. E. Clearance of 10 nM [^35^S]MMP-1 after 24 h incubation with HTB94 cells was compared to that of 10 nM [^35^S]TIMP-3, [^35^S]MMP-1/TIMP-3, [^35^S]MMP-1/TIMP-1 and [^35^S]MMP-1/TIMP-2. Values for [^35^S]MMP-1, [^35^S]MMP-1/TIMP-3, [^35^S]MMP-1/TIMP-1 and [^35^S]MMP-1/TIMP-2 were extrapolated from panel A-D an represented, together with those for [^35^S]TIMP-3 clearance, as mean values of intact protein ([^35^S]MMP-1 or [^35^S]TIMP-3 remained in the media □±□ standard deviation, and analysed by one-way ANOVA, followed by Dunnett’s multiple comparisons test where [^35^S]MMP-1 was taken as a control; ns non significant, *p□<□ 0.05, **p□<□ 0.01, ***p□<□ 0.005; n=3-5). F. Sensograms from a SPR analysis showing binding of 10 nM MMP-1/TIMP-3 (black solid line), MMP-1/TIMP-1 (black dashed line) or MMP-1/TIMP-2 (grey dashed line) to LRP-1 immobilised on a sensor chip.

In conclusion, we identified MMP-1 as a novel LRP-1 ligand that, similarly to other metalloproteinases, is actively endocytosed and degraded by cells. TIMP-3, but not TIMP-1 or TIMP-2, increased MMP-1 internalisation rate when in complex with the proteinase.

## DISCUSSION

Two classes of metalloproteinases, MMPs and ADAMTSs, play a central role in ECM turnover, together with the ADAMs that are majorly involved in ectodomain shedding of cell-surface components. The activity of these enzymes is finely regulated by their endogenous inhibitors, known as TIMPs. LRP-1 is an endocytic receptor that has been reported to mediate the internalisation and degradation of a number of MMPs, ADAMTSs and TIMPs, thus functioning as a master regulator of ECM turnover (9,10,12,14-16,18). Levels of these proteins in the ECM can tightly correlate to their affinity for the receptor. For example, TIMP-3, which was reported to bind to LRP-1 with high affinity, is hardly found extracellularly unless its interaction with LRP-1 and subsequent endocytosis are prevented (18,38). Investigating affinities for LRP-1 of different metalloproteinases, TIMPs and metalloproteinase/TIMP complexes can provide useful information about turnover of these proteins and potential effects on ECM turnover.

In the current study, we used SPR to analyse in a systematic manner binding to LRP-1 of a number of MMPs, ADAMs, ADAMTSs, TIMPs and metalloproteinase/TIMP complexes. We found that, among these proteins, TIMP-3 bound to LRP-1 with highest affinity. Furthermore, TIMP-3 increases the affinity for LRP-1 of target proteinases that bind to LRP-1 by themselves, such as MMP-1, MMP-13 and ADAMTS-4, but also that of metalloproteinases, including MMP-14, ADAM10 and ADAM17, which have no affinity for the receptor. ADAMTS-5 binds to and is rapidly endocytosed by LRP-1 (37), and we found that the affinity of ADAMTS-5 lacking the C-terminal thrombospondin domain (ADAMTS5-2) for the receptor was similar to that of TIMP-3. After secretion, ADAMTS-5 is extracellularly processed to truncated forms that acquire distinct features from the full-length counterpart (39,40), including different LRP-1 affinities and endocytosis rates (14). Expression and purification of recombinant full-length ADAMTS-5 is inefficient, so we did not analyse the full-length enzyme in this study. While TIMP-3 had little effects on the affinity of ADAMTS-5 for LRP-1, it doubled the affinity of a truncated form of ADAMTS-5 lacking of the cysteine-rich and spacer domain (ADAMTS5-4) for the receptor, suggesting TIMP-3 may promote scavenging of smaller, degraded forms of the enzyme.

Similarly to TIMP-3, TIMP-1 and TIMP-2 directly bound to LRP-1, although with lower affinities. Both TIMP-1 and TIMP-2 increased the affinities of MMPs for LRP-1, but not to the same extent as TIMP-3. This indicates that there are at least two modes by which inhibitors can increase proteinase affinity for LRP-1. In the simplest case, binding may be mediated primarily through residues on the inhibitor, such that the inhibitor/proteinase complex has similar affinity for LRP-1 as the free inhibitor alone. Such behaviour was observed for TIMP-3, with TIMP-3/MMP-13, TIMP-3/MMP-1 and TIMP-3/MMP-14 complexes having similar affinity for LRP-1 as free TIMP-3. We demonstrated that this mode of binding is able to support biologically relevant bridging of the MMP to LRP-1, as TIMP-3 was able to promote MMP-1 endocytosis by HTB94 chondrosarcoma cells.

Alternatively, the proteinase-inhibitor complex may bind with higher affinity than either protein alone. We observed that TIMP-1 and TIMP-2 complexes with both MMP-1 and MMP-13 had ∼2-5-fold higher affinity for LRP-1 than any of the four proteins alone. This is in line with what has been reported for the proMMP-2/TIMP-2 complex, which has ∼10-fold higher affinity for LRP-1 than either component alone (9). This may be due to a conformational change induced by proteinase-inhibitor interaction, as is the case with α_2_-macroglobulin, which has low affinity for LRP-1 in its free form, but which undergoes a conformational change upon proteinase binding that increases affinity of the complex for LRP-1. Alternatively, both enzyme and inhibitor may directly participate in LRP-1 binding, each contributing a lysine residue to form the minimal LRP-1 binding unit (32). We consider this the more likely mechanism, as TIMPs are considered to be conformationally stable, with only minor structural rearrangements upon interaction with MMPs. However, this mode of binding appears to be insufficient to support bridging to LRP-1 in a complex biological environment, as we found that neither TIMP-1 nor TIMP-2 could increase the rate of MMP-1 endocytosis by HTB94 chondrosarcoma cells. Thus, our kinetic data indicate that TIMP-3 bridges MMP binding to LRP-1 though a different mechanism to that of TIMP-1 and TIMP-2, and that only TIMP-3 can promote scavenging of these target enzymes from the extracellular environment.

The observation that TIMP-3/metalloproteinase complexes and free TIMP-3 have similar affinity for LRP-1 to that of TIMP-3 alone suggests that TIMP-3 interacts with LRP-1 through a site distinct from its inhibitory ridge. Studies of a number of LRP-1 ligands suggest that a pair of basic residues is responsible for binding to acidic pockets on the complement-type repeats (CRs) of the receptor (41,42). For example, extensive mutagenesis and crystallography studies indicate that RAP binds to LRP-1 via two lysine residues, K256 and K270, which are located 22 Å apart on the surface of the molecule (32). TIMP-3 contains a cluster of lysine and arginine residues situated on the opposite side to the inhibitory ridge. This basic region mediates binding to sulfated proteoglycans of the extracellular matrix (43). We previously found that heparin blocks TIMP-3 binding to LRP-1 (30), suggesting that the heparin-binding and LRP-1-binding sites of TIMP-3 overlap, and supporting the postulate that the metalloproteinase-inhibitory region of TIMP-3 is spatially separated from the LRP-1-binding and overlapping ECM-binding basic region.

There is emerging evidence that metalloproteinases have functions other than proteolytic cleavage, and that these functions are often related to their non-catalytic domains. For example, a non-catalytic C-terminal fragment of MMP-12 has been shown to have antimicrobial activity (44). ADAMTS-1 and ADAMTS-4 display anti-angiogenic properties through their ability to bind VEGF and sequester it from VEGFR (45,46). Fragments of ADAMTS-5 derived from its ancillary domains have been shown to be anti-tumorigenic both *in vitro* and *in vivo* (47). ADAMTS-12 modulates neutrophil apoptosis, likely by interacting with CD36 and CD47 through its thrombospondin domains (48). Although it has not been explicitly demonstrated, it is likely that at least some of these non-catalytic functions are maintained when metalloproteinases interact with TIMPs. In addition to its role in inhibiting the proteolytic activity of target metalloproteinases, TIMP-3 may thus also inhibit non-proteolytic activities of target proteinases by facilitating their LRP-1-mediated endocytosis. This would position TIMP-3 as a central regulator of metalloproteinase levels and activity in the extracellular environment.

It is interesting to speculate that competition between different TIMPs for metalloproteinases could lead to diverse biological responses. TIMP-1 and TIMP-2 would serve primarily to inhibit the catalytic activity of the proteinases, but would not facilitate their endocytic clearance. TIMP-3, on the other hand, is uniquely able to address the enzymes to intracellular degradation and therefore regulate both their proteolytic and non-proteolytic functions. We found that TIMP-3 also bridges binding of members of the ADAM and ADAMTS families to LRP-1. The ectodomains of ADAM10 and ADAM17 do not interact with the receptor unless they are in complex with TIMP-3. This may explain why TIMP-3 decreases cell surface levels of ADAM10 (49).

In conclusion, this work shows that TIMP-3 can promote LRP-1-mediated endocytosis of target metalloproteinases by facilitating their binding to LRP-1. As such, TIMP-3 can inhibit both proteolytic-dependent and - independent activities of metalloproteinases in the extracellular environment.

## REFERENCES

1. Nagase, H., and Woessner, J. F., Jr. (1999) Matrix metalloproteinases. J. Biol. Chem. 274, 21491–21494

2. Apte, S. S. (2009) A disintegrin-like and metalloprotease (reprolysin-type) with thrombospondin type 1 motif (ADAMTS) superfamily: functions and mechanisms. J Biol Chem 284, 31493–31497

3. Seals, D. F., and Courtneidge, S. A. (2003) The ADAMs family of metalloproteases: multidomain proteins with multiple functions. Genes Dev 17, 7–30

4. Brew, K., and Nagase, H. (2010) The tissue inhibitors of metalloproteinases (TIMPs): an ancient family with structural and functional diversity. Biochim Biophys Acta 1803, 55–71

5. Murphy, G., and Nagase, H. (2008) Reappraising metalloproteinases in rheumatoid arthritis and osteoarthritis: destruction or repair? Nat. Clin. Pract. Rheumatol. 4, 128–135

6. Nagase, H., Visse, R., and Murphy, G. (2006) Structure and function of matrix metalloproteinases and TIMPs. Cardiovasc Res 69, 562–573

7. Egeblad, M., and Werb, Z. (2002) New functions for the matrix metalloproteinases in cancer progression. Nat Rev Cancer 2, 161–174

8. Nagase, H., Visse, R., and Murphy, G. (2006) Structure and function of matrix metalloproteinases and TIMPs. Cardiovasc Res 69, 562–573

9. Emonard, H., Bellon, G., Troeberg, L., Berton, A., Robinet, A., Henriet, P., Marbaix, E., Kirkegaard, K., Patthy, L., Eeckhout, Y., Nagase, H., Hornebeck, W., and Courtoy, P. J. (2004) Low density lipoprotein receptor-related protein mediates endocytic clearance of pro-MMP-2.TIMP-2 complex through a thrombospondin-independent mechanism. J Biol Chem 279, 54944–54951

10. Hahn-Dantona, E., Ruiz, J. F., Bornstein, P., and Strickland, D. K. (2001) The low density lipoprotein receptor-related protein modulates levels of matrix metalloproteinase 9 (MMP-9) by mediating its cellular catabolism. J. Biol. Chem. 276, 15498–15503

11. Barmina, O. Y., Walling, H. W., Fiacco, G. J., Freije, J. M., Lopez-Otin, C., Jeffrey, J. J., and Partridge, N. C. (1999) Collagenase-3 binds to a specific receptor and requires the low density lipoprotein receptor-related protein for internalization. J. Biol. Chem. 274, 30087–30093

12. Yamamoto, K., Okano, H., Miyagawa, W., Visse, R., Shitomi, Y., Santamaria, S., Dudhia, J., Troeberg, L., Strickland, D. K., Hirohata, S., and Nagase, H. (2016) MMP-13 is constitutively produced in human chondrocytes and co-endocytosed with ADAMTS-5 and TIMP-3 by the endocytic receptor LRP1. Matrix Biol

13. Scilabra, S. D., Pigoni, M., Pravata, V., Schatzl, T., Muller, S. A., Troeberg, L., and Lichtenthaler, S. F. (2018) Increased TIMP-3 expression alters the cellular secretome through dual inhibition of the metalloprotease ADAM10 and ligand-binding of the LRP-1 receptor. Sci Rep 8, 14697

14. Yamamoto, K., Owen, K., Parker, A. E., Scilabra, S. D., Dudhia, J., Strickland, D. K., Troeberg, L., and Nagase, H. (2014) Low density lipoprotein receptor-related protein 1 (LRP1)-mediated endocytic clearance of a disintegrin and metalloproteinase with thrombospondin motifs-4 (ADAMTS-4): functional differences of non-catalytic domains of ADAMTS-4 and ADAMTS-5 in LRP1 binding. J Biol Chem 289, 6462–6474

15. Yamamoto, K., Troeberg, L., Scilabra, S. D., Pelosi, M., Murphy, C. L., Strickland, D. K., and Nagase, H. (2013) LRP-1-mediated endocytosis regulates extracellular activity of ADAMTS-5 in articular cartilage. FASEB J 27, 511–521

16. Thevenard, J., Verzeaux, L., Devy, J., Etique, N., Jeanne, A., Schneider, C., Hachet, C., Ferracci, G., David, M., Martiny, L., Charpentier, E., Khrestchatisky, M., Rivera, S., Dedieu, S., and Emonard, H. (2014) Low-density lipoprotein receptor-related protein-1 mediates endocytic clearance of tissue inhibitor of metalloproteinases-1 and promotes its cytokine-like activities. PLoS One 9, e103839

17. Troeberg, L., Fushimi, K., Khokha, R., Emonard, H., Ghosh, P., and Nagase, H. (2008) Calcium pentosan polysulfate is a multifaceted exosite inhibitor of aggrecanases. FASEB J. 22, 3515–3524

18. Scilabra, S. D., Troeberg, L., Yamamoto, K., Emonard, H., Thogersen, I., Enghild, J. J., Strickland, D. K., and Nagase, H. (2013) Differential regulation of extracellular tissue inhibitor of metalloproteinases-3 levels by cell membrane-bound and shed low density lipoprotein receptor-related protein 1. J Biol Chem 288, 332–342

19. Hahn-Dantona, E., Ruiz, J. F., Bornstein, P., and Strickland, D. K. (2001) The low density lipoprotein receptor-related protein modulates levels of matrix metalloproteinase 9 (MMP-9) by mediating its cellular catabolism. J Biol Chem 276, 15498–15503

20. Emonard, H., Bellon, G., Troeberg, L., Berton, A., Robinet, A., Henriet, P., Marbaix, E., Kirkegaard, K., Patthy, L., Eeckhout, Y., Nagase, H., Hornebeck, W., and Courtoy, P. J. (2004) Low density lipoprotein receptor-related protein mediates endocytic clearance of pro-MMP-2.TIMP-2 complex through a thrombospondin-independent mechanism. J Biol Chem 279, 54944–54951

21. Chung, L., Dinakarpandian, D., Yoshida, N., Lauer-Fields, J. L., Fields, G. B., Visse, R., and Nagase, H. (2004) Collagenase unwinds triple-helical collagen prior to peptide bond hydrolysis. EMBO J 23, 3020–3030

22. Kashiwagi, M., Enghild, J. J., Gendron, C., Hughes, C., Caterson, B., Itoh, Y., and Nagase, H. (2004) Altered proteolytic activities of ADAMTS-4 expressed by C-terminal processing. J. Biol. Chem. 279, 10109–10119

23. Gendron, C., Kashiwagi, M., Lim, N. H., Enghild, J. J., Thogersen, I. B., Hughes, C., Caterson, B., and Nagase, H. (2007) Proteolytic activities of human ADAMTS-5: comparative studies with ADAMTS-4. J Biol Chem 282, 18294–18306

24. Troeberg, L., Fushimi, K., Scilabra, S. D., Nakamura, H., Dive, V., Thogersen, I. B., Enghild, J. J., and Nagase, H. (2009) The C-terminal domains of ADAMTS-4 and ADAMTS-5 promote association with N-TIMP-3. Matrix Biol. 28, 463–469

25. Troeberg, L., Tanaka, M., Wait, R., Shi, Y. E., Brew, K., and Nagase, H. (2002) E. coli expression of TIMP-4 and comparative kinetic studies with TIMP-1 and TIMP-2: insights into the interactions of TIMPs and matrix metalloproteinase 2 (gelatinase A). Biochemistry 41, 15025–15035

26. Suzuki, K., Kan, C. C., Hung, W., Gehring, M. R., Brew, K., and Nagase, H. (1998) Expression of human pro-matrix metalloproteinase 3 that lacks the N-terminal 34 residues in Escherichia coli: autoactivation and interaction with tissue inhibitor of metalloproteinase 1 (TIMP-1). Biol Chem 379, 185–191

27. O’Connell, J. P., Willenbrock, F., Docherty, A. J., Eaton, D., and Murphy, G. (1994) Analysis of the role of the COOH-terminal domain in the activation, proteolytic activity, and tissue inhibitor of metalloproteinase interactions of gelatinase B. J Biol Chem 269, 14967–14973

28. Stura, E. A., Visse, R., Cuniasse, P., Dive, V., and Nagase, H. (2013) Crystal structure of full-length human collagenase 3 (MMP-13) with peptides in the active site defines exosites in the catalytic domain. FASEB J 27, 4395–4405

29. Huang, W., Suzuki, K., Nagase, H., Arumugam, S., Van Doren, S. R., and Brew, K. (1996) Folding and characterization of the amino-terminal domain of human tissue inhibitor of metalloproteinases-1 (TIMP-1) expressed at high yield in E. coli. FEBS Lett 384, 155–161

30. Scilabra, S. D., Troeberg, L., Yamamoto, K., Emonard, H., Thogersen, I., Enghild, J. J., Strickland, D. K., and Nagase, H. (2013) Differential regulation of extracellular tissue inhibitor of metalloproteinases-3 levels by cell membrane-bound and shed low density lipoprotein receptor-related protein 1. J Biol Chem 288, 332–342

31. Chung, L., Shimokawa, K., Dinakarpandian, D., Grams, F., Fields, G. B., and Nagase, H. (2000) Identification of the (183)RWTNNFREY(191) region as a critical segment of matrix metalloproteinase 1 for the expression of collagenolytic activity. J Biol Chem 275, 29610–29617

32. van den Biggelaar, M., Sellink, E., Klein Gebbinck, J. W., Mertens, K., and Meijer, A. B. (2011) A single lysine of the two-lysine recognition motif of the D3 domain of receptor-associated protein is sufficient to mediate endocytosis by low-density lipoprotein receptor-related protein. Int J Biochem Cell Biol 43, 431–440

33. Migliorini, M. M., Behre, E. H., Brew, S., Ingham, K. C., and Strickland, D. K. (2003) Allosteric modulation of ligand binding to low density lipoprotein receptor-related protein by the receptor-associated protein requires critical lysine residues within its carboxyl-terminal domain. J Biol Chem 278, 17986–17992

34. Lee, D., Walsh, J. D., Mikhailenko, I., Yu, P., Migliorini, M., Wu, Y., Krueger, S., Curtis, J. E., Harris, B., Lockett, S., Blacklow, S. C., Strickland, D. K., and Wang, Y. X. (2006) RAP uses a histidine switch to regulate its interaction with LRP in the ER and Golgi. Mol Cell 22, 423–430

35. Mantuano, E., Inoue, G., Li, X., Takahashi, K., Gaultier, A., Gonias, S. L., and Campana, W. M. (2008) The hemopexin domain of matrix metalloproteinase-9 activates cell signaling and promotes migration of schwann cells by binding to low-density lipoprotein receptor-related protein. J Neurosci 28, 11571–11582

36. Zeng, S., Yu, X., Law, W. C., Zhang, Y., Hu, R., Dinh, X. Q., Ho, H. P., and Yong, K. T. (2013) Size dependence of Au NP-enhanced surface plasmon resonance based on differential phase measurement. Sensors and Actuators B: Chemical 176, 1128–1133

37. Yamamoto, K., Troeberg, L., Scilabra, S. D., Pelosi, M., Murphy, C. L., Strickland, D. K., and Nagase, H. (2013) LRP-1-mediated endocytosis regulates extracellular activity of ADAMTS-5 in articular cartilage. Faseb J 27, 511–521

38. Scilabra, S. D., Yamamoto, K., Pigoni, M., Sakamoto, K., Muller, S. A., Papadopoulou, A., Lichtenthaler, S. F., Troeberg, L., Nagase, H., and Kadomatsu, K. (2017) Dissecting the interaction between tissue inhibitor of metalloproteinases-3 (TIMP-3) and low density lipoprotein receptor-related protein-1 (LRP-1): Development of a “TRAP” to increase levels of TIMP-3 in the tissue. Matrix Biol 59, 69–79

39. Gao, G., Westling, J., Thompson, V. P., Howell, T. D., Gottschall, P. E., and Sandy, J. D. (2002) Activation of the proteolytic activity of ADAMTS4 (aggrecanase-1) by C-terminal truncation. J Biol Chem 277, 11034–11041

40. Plaas, A., Osborn, B., Yoshihara, Y., Bai, Y., Bloom, T., Nelson, F., Mikecz, K., and Sandy, J. D. (2007) Aggrecanolysis in human osteoarthritis: confocal localization and biochemical characterization of ADAMTS5-hyaluronan complexes in articular cartilages. Osteoarthritis Cartilage 15, 719–734

41. Lillis, A. P., Mikhailenko, I., and Strickland, D. K. (2005) Beyond endocytosis: LRP function in cell migration, proliferation and vascular permeability. J Thromb Haemost 3, 1884–1893

42. Andersen, O. M., Christensen, L. L., Christensen, P. A., Sorensen, E. S., Jacobsen, C., Moestrup, S. K., Etzerodt, M., and Thogersen, H. C. (2000) Identification of the minimal functional unit in the low density lipoprotein receptor-related protein for binding the receptor-associated protein (RAP). A conserved acidic residue in the complement-type repeats is important for recognition of RAP. J Biol Chem 275, 21017–21024

43. Lee, M. H., Atkinson, S., and Murphy, G. (2007) Identification of the extracellular matrix (ECM) binding motifs of tissue inhibitor of metalloproteinases (TIMP)-3 and effective transfer to TIMP-1. J Biol Chem 282, 6887–6898

44. Marchant, D. J., Bellac, C. L., Moraes, T. J., Wadsworth, S. J., Dufour, A., Butler, G. S., Bilawchuk, L. M., Hendry, R. G., Robertson, A. G., Cheung, C. T., Ng, J., Ang, L., Luo, Z., Heilbron, K., Norris, M. J., Duan, W., Bucyk, T., Karpov, A., Devel, L., Georgiadis, D., Hegele, R. G., Luo, H., Granville, D. J., Dive, V., McManus, B. M., and Overall, C. M. (2014) A new transcriptional role for matrix metalloproteinase-12 in antiviral immunity. Nat Med 20, 493–502

45. Hsu, Y. P., Staton, C. A., Cross, N., and Buttle, D. J. (2012) Antiangiogenic properties of ADAMTS-4 in vitro. Int J Exp Pathol 93, 70–77

46. Luque, A., Carpizo, D. R., and Iruela-Arispe, M. L. (2003) ADAMTS1/METH1 inhibits endothelial cell proliferation by direct binding and sequestration of VEGF165. J Biol Chem 278, 23656–23665

47. Kumar, S., Sharghi-Namini, S., Rao, N., and Ge, R. (2012) ADAMTS5 functions as an anti-angiogenic and anti-tumorigenic protein independent of its proteoglycanase activity. Am J Pathol 181, 1056–1068

48. Moncada-Pazos, A., Obaya, A. J., Llamazares, M., Heljasvaara, R., Suarez, M. F., Colado, E., Noel, A., Cal, S., and Lopez-Otin, C. (2012) ADAMTS-12 metalloprotease is necessary for normal inflammatory response. J Biol Chem 287, 39554–39563

49. Hoe, H. S., Cooper, M. J., Burns, M. P., Lewis, P. A., van der Brug, M., Chakraborty, G., Cartagena, C. M., Pak, D. T., Cookson, M. R., and Rebeck, G. W. (2007) The metalloprotease inhibitor TIMP-3 regulates amyloid precursor protein and apolipoprotein E receptor proteolysis. J Neurosci 27, 10895–10905

